# Reducing background ion burden in tributylamine ion-pairing LC-MS improves signal intensity and feature coverage in metabolomics

**DOI:** 10.64898/2026.06.24.734057

**Authors:** Anna R. Tarach, Michael P. Vincent, Abigail E. Ellis, Christine Isaguirre, Amy Caudy, Ryan D. Sheldon

## Abstract

Background chemical ions are a pervasive but often underappreciated limitation in LC-MS metabolomics, where they can suppress analyte signal, obscure endogenous metabolites, increase spectral complexity, and consume MS/MS acquisition events. Tributylamine (TBA) ion-pairing reversed-phase LC-MS provides stable retention and broad coverage of polar anionic metabolites, including central carbon intermediates, nucleotides, cofactors, and bile acids, but the background burden introduced by the ion-pairing reagent itself has not been systematically addressed. Here, we identify commercial TBA as a major source of nonbiological contaminant ions and develop a practical strategy to reduce background burden while preserving metabolite coverage. Serial solid-phase extraction of TBA using orthogonal reversed-phase, strong anion-exchange, and strong cation-exchange sorbents removed chemically diverse contaminants, including isobaric background ions that interfered with endogenous hydroxybutyrate isomers. We further optimized the workflow by reducing medronic acid concentration, restricting medronic acid to the organic mobile phase, replacing phosphoric-acid column conditioning with metal-passivated column hardware, and adding EDTA to the sample reconstitution solvent to improve citrate detection. In mouse liver extracts, the optimized method increased signal intensity for most annotated metabolites and improved the fraction of full-scan ion current attributable to target analytes. Method optimization also altered compound-specific retention behavior, resolving some co-elution-based interferences while introducing new suppression relationships for selected analytes. Across mouse liver, human B lymphocytes, and NIST SRM 1950 plasma, the optimized workflow increased total feature detection by 45%, 72%, and 42%, respectively, and improved the number of low-variance features, precursors with data-dependent MS/MS spectra, and MS/MS library matches. These findings establish background-ion mitigation as a central design principle for LC-MS method development. More broadly, this work provides a generalizable framework for identifying, reducing, and validating reagent- and additive-derived background to improve targeted and untargeted LC-MS data quality.

## Introduction

Liquid chromatography–mass spectrometry is widely used for metabolomics because it can detect large numbers of chemically diverse metabolites spanning large concentration ranges in complex biological matrices. However, the breadth of the metabolome also creates a persistent analytical challenge because metabolites differ widely in polarity, charge state, abundance, chemical stability, and ionization efficiency. As a result, the chromatographic method is not simply a front-end separation step, but a major determinant of metabolite coverage, signal intensity, reproducibility, and annotation confidence^1, 2^.

Polar and anionic metabolites remain difficult to measure by LC–MS because many are poorly retained by conventional reversed-phase chromatography. This group includes central carbon metabolites such as glycolytic intermediates, tricarboxylic acid cycle intermediates, sugar phosphates, nucleotides, and related cofactors. Hydrophilic interaction liquid chromatography (HILIC) is commonly used for these analytes, but HILIC methods can be sensitive to column equilibration, retention-time drift, and method-to-method variability, which can limit the use of retention time for metabolite annotation^3^. Reversed-phase ion-pairing chromatography offers a complementary strategy by using volatile amine additives to retain anionic metabolites on hydrophobic stationary phases. Tributylamine (TBA)-based methods have been widely used for targeted and untargeted analysis of polar central carbon metabolites, nucleotides, and cofactors^4–6^. Because these separations are performed on reversed-phase columns, they can also retain later-eluting amphipathic metabolites, such as bile acids, thereby extending coverage beyond the most polar portion of metabolism. Thus, TBA ion-pairing provides a stable, broad-coverage approach that complements HILIC-based metabolomics workflows.

Despite its advantages, TBA ion-pairing introduces background-related challenges that may arise from the same chemistry that makes the method useful. Under acidic mobile-phase conditions, TBA is protonated to tributylammonium, which promotes retention of anionic metabolites on reversed-phase stationary phases. This anion-binding capacity may also make low-background TBA difficult to source, as illustrated by the use of tributylamine strong-base anion-exchange resins for nitrate removal in water-treatment applications^7^. More generally, ion-pairing reagents and mobile-phase additives can contribute chemical background and ion suppression^8^, and TBA is known to cause strong positive-mode ion suppression, generally limiting TBA-based metabolomics to negative-ion acquisition^5^. However, the extent to which TBA-derived background impairs negative-mode metabolomics through ion suppression, isobaric interference, increased spectral complexity, and nonbiological data-dependent MS/MS acquisition remains poorly defined.

A second challenge is the susceptibility of polar anionic metabolites to adsorption on metal or metal-oxide surfaces in the LC flow path. Phosphorylated metabolites and multicarboxylate organic acids can show poor recovery, peak tailing, carryover, and reduced signal because of interactions with stainless steel tubing, frits, column hardware, and other exposed metal components^9, 10^. Medronic acid and metal-passivated LC hardware can mitigate these losses and improve detection of metal-sensitive analytes^9–11^. However, these additives like medronic acid can add background signal or suppress ionization, while phosphoric-acid column conditioning is labor-intensive and may lose effectiveness over time.

Here, we report an optimized low-background TBA ion-pairing LC-MS workflow for improved metabolomics data quality. We first identify TBA as a major source of background contaminant ions and show that these contaminants can be reduced by serial solid-phase extraction using orthogonal sorbent chemistries. We then optimize metal-mitigation strategies by minimizing medronic acid burden, replacing phosphoric-acid column conditioning with metal-passivated column hardware, and adding EDTA to the sample reconstitution solvent to improve citrate detection. Finally, we benchmark the optimized method against the initial conditions across mouse liver, human B lymphocyte, and human plasma extracts. The optimized workflow increases signal intensity for shared features, expands feature detection, improves acquisition of interpretable MS/MS spectra, and increases MS/MS library matches across diverse biological matrices.

## Results and Discussion

Tributylamine-based ion-pairing chromatography provides excellent retention and separation of polar metabolites involved in central carbon metabolism and related pathways, including nucleotides, amino acids, tricarboxylic acid cycle intermediates, sugar phosphates, and other polar metabolites (Fig 1A-B). With proper maintenance, the method also provides long-term stability in peak shape and retention time, making it an attractive alternative to HILIC, which is often prone to retention-time drift and frequent column replacement. As evidence of this stability, using system suitability data from our lab, we demonstrate minimal variation in retention time over a 400-day period encompassing more than 3,000 sample injections and four column changes (Fig 1C). However, the method produces a relatively high background signal from contaminant ions of unknown origin, leading to both ion suppression and isobaric interference with endogenous metabolites. We hypothesized that many of these contaminants originate from the tributylamine additive, which is not available as a high-purity LC-MS-grade reagent and by its nature as a protonatable tertiary primary amine is prone to binding anions during manufacturing and processing.

**Figure 1.**
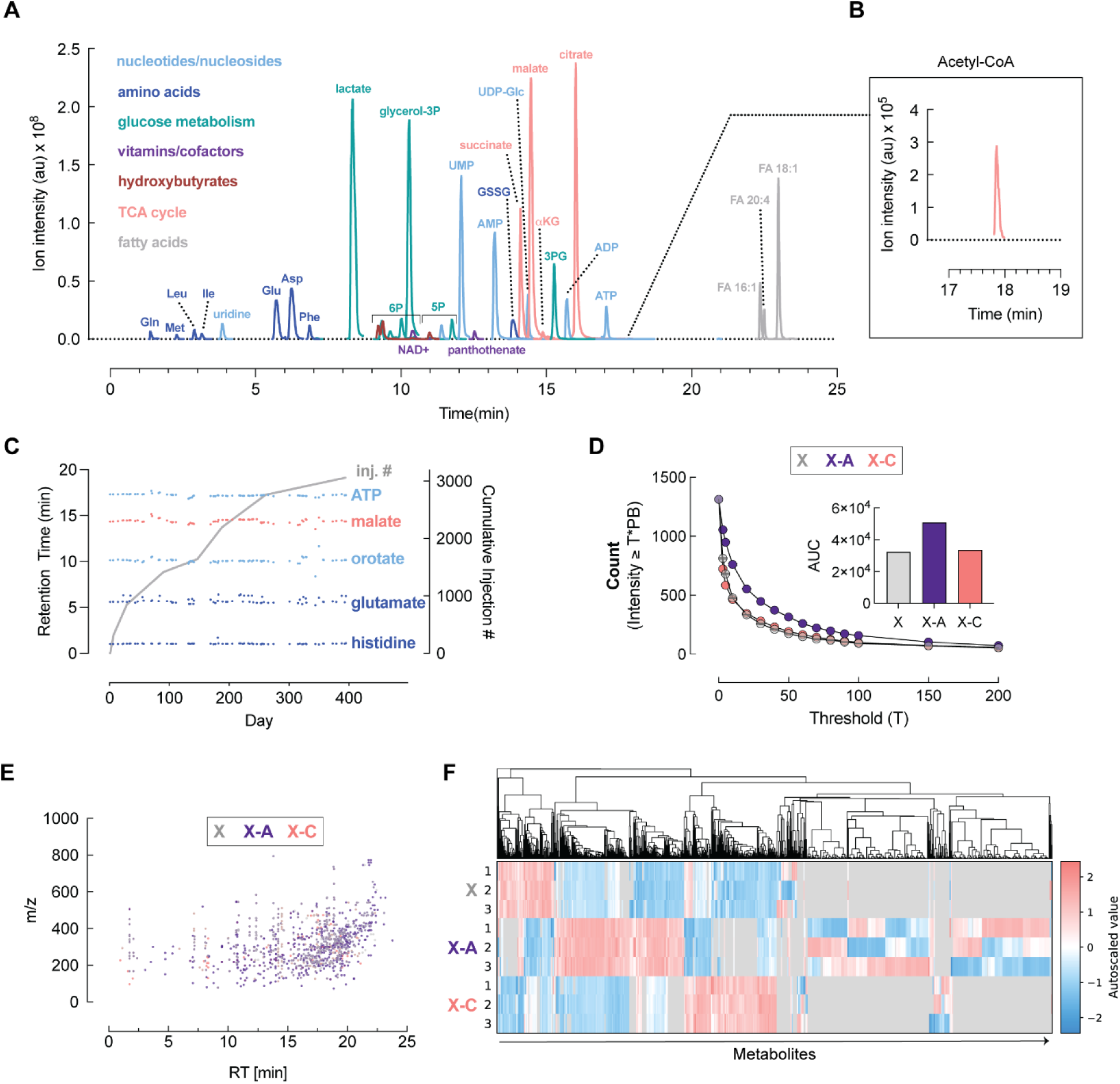
Tributylamine ion-paired LC-MS offers long-term stability and good coverage of central carbon metabolites but suffers from high contaminate load. **(A)** Overlaid representative extracted ion chromatogram of select central carbon metabolites using IP LC-MS on mouse liver metabolite extract. **(B)** Inset acetyl-CoA extracted ion chromatogram. **(C)** System suitability mixture of histidine, glutamate, orotate, malate, and ATP injected over a 400-day period **(D)** Features detected in response to increased background thresholds. Tributylamine was purified using Strata X, Strata X-A, and Strata X-C columns. Counts represent the number of features having a level that exceeds the level observed in the process blank (PB) multiplied by the specified threshold (T). Inset: integrated area under curve (AUC) from each sorbent type. **(E)** Mass-to-charge (m/z) and retention time (RT) profile of features observed with greater than 10-fold signal over blank following filtration with Strata X, Stata X-A, or Strata X-C columns. **(F)** Heatmap of contaminants removed from TBA by Strata X, Strata X-A, and Strata X-C columns and analyzed by IP LC-MS. 1568 features are shown with peak area >10^6^ and at least 10X over blank. Presented as auto-scaled Z score. Experiments in **D-F** used n=3 technical replicates per condition.

To test this hypothesis, we subjected tributylamine to serial solid-phase extraction using reversed-phase polymeric and ion-exchange sorbents to remove a broad range of contaminants. Specifically, tributylamine was passed sequentially through Strata-X, Strata-X-A, and Strata-X-C cartridges. To evaluate contaminant capture by each sorbent, we eluted compounds retained on the cartridges after tributylamine purification and analyzed these eluates by LC-MS. Matched process blanks were prepared using identically handled cartridges that received LC-MS-grade water instead of tributylamine. Total ion chromatograms showed many additional peaks in eluates from tributylamine-treated cartridges compared with water blanks (Figure 1D,E). Untargeted feature detection identified 1,314 features across the chromatographic run that were present in eluates from at least one sorbent, had a peak rating greater than 7 (Compound Discoverer v3.5), had an ion intensity greater than 10, and were enriched at least 10-fold relative to process blanks (Figure E). Strata-X cartridges removed the largest number of contaminants, whereas Strata-X-A and Strata-X-C captured distinct additional subsets of contaminants (Figure 1F).

These results indicate that tributylamine is a major source of background contaminants, many of which can be depleted by serial SPE using orthogonal sorbent chemistries. The chemical diversity of the removed features is consistent with the intended function of TBA as an ion-pairing reagent: under acidic mobile-phase conditions, TBA is protonated to tributylammonium, which promotes retention of anionic compounds on reversed-phase stationary phases. This same anion-binding chemistry may also make low-background TBA difficult to source or produce, because protonated or immobilized tributylamine chemistries can associate with diverse anionic species, as illustrated by the use of tributylamine-derived strong-base anion-exchange materials for nitrate removal in water-treatment applications^7^. Mobile-phase-derived background is a recognized source of interference in gradient LC–MS: trace impurities in mobile-phase water can accumulate on the column during equilibration, elute during the gradient, and suppress or enhance analyte signals^12^. Common negative-mode chemical noise can also arise from mobile-phase components, additives, and contaminant clusters^13^. Sorbent-based background mitigation has precedent in LC–MS/MS workflows, where trap columns have been used to retain and delay mobile-phase-derived contaminants, thereby separating background ions from sample-derived analyte signals^14^. Here, we extend this principle upstream by applying serial orthogonal SPE directly to commercial TBA before mobile-phase preparation, reducing reagent-derived background ions before they enter the LC–MS workflow.

Among the features depleted by TBA purification, 603 had at least one accurate mass match withing 5ppm of a ChemSpider annotated compound, indicating that TBA-derived contaminants may be a substantial source of isobaric interference. One of the highest-abundance contaminants removed was a feature at *m/z* 103.0401. This is consistent with the deprotonated ion [M-H]^-^ of short-chain hydroxycarboxylic acids, which includes common biologically occurring structural isomers 2-hydroxybutyrate, 3-hydroxybutyrate, 3-hydroxyisobutyrate, and 4-hydroxybutyrate. We compared the extracted ion chromatograms of m/z 103.0401 of water-blank injections using mobile phases prepared with unpurified or SPE purified TBA. Mobile phases containing unpurified TBA produced a high background signal for this ion across the chromatogram, whereas this signal was markedly reduced when SPE-purified TBA was used (Figure 2A). In liver metabolite extracts, this high background obscured the resolution of endogenous hydroxybutyrate isomers (Figure 2B). In contrast, SPE-purified TBA reduced the *m/z* 103.0401 background sufficiently to chromatographically resolve 3-hydroxyisobutyrate, 3-hydroxybutyrate, 2-hydroxybutyrate, and 4-hydroxybutyrate (Figure 2C).

**Figure 2.**
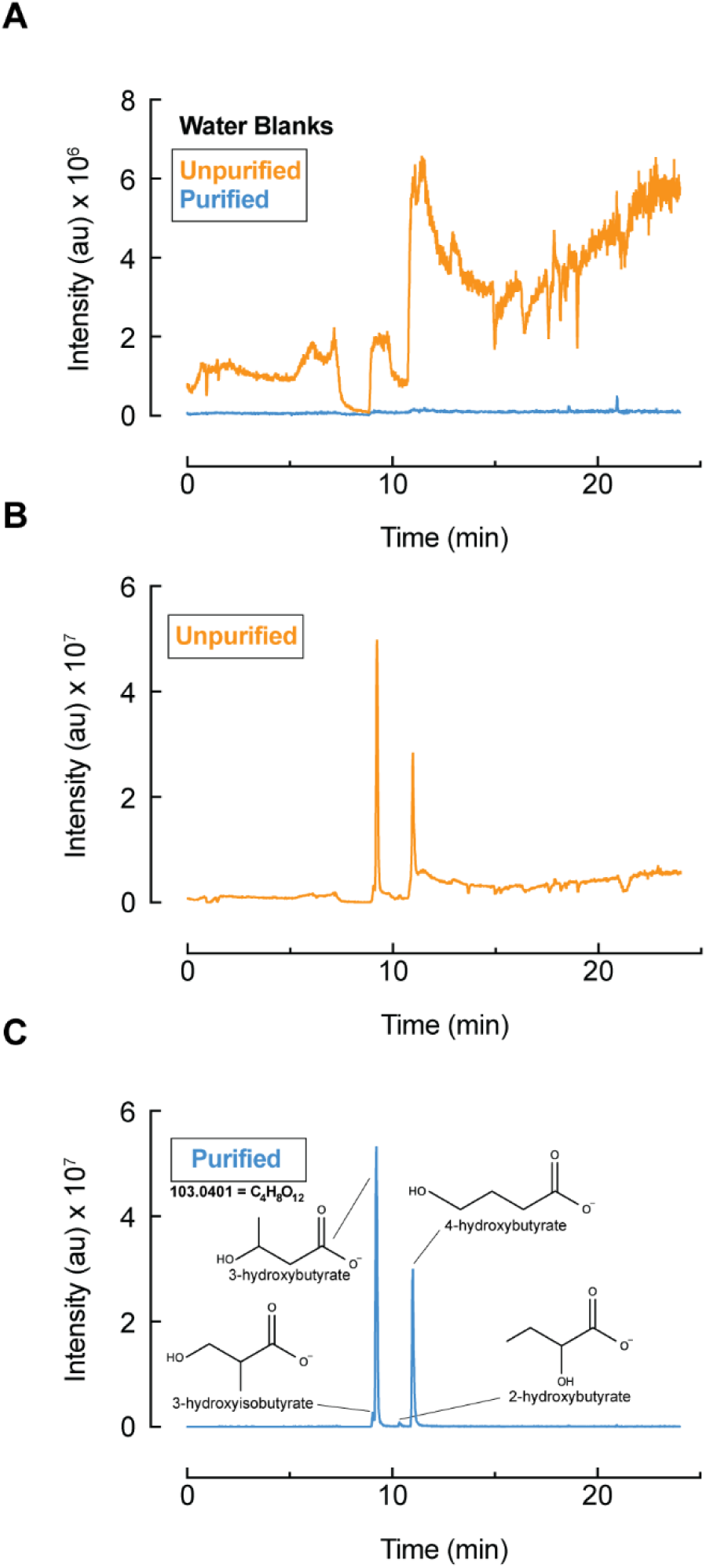
TBA purification removes isobaric inference that allows for resolution of biologically occurring hydroxybutyrates. **(A)** Representative extracted ion chromatogram of [103.0401]^-^ in water blank samples using un-purified or purified TBA in the mobile phases**. (B)** Representative extracted ion chromatogram of [103.0401]^-^ in mouse liver metabolite extracts using unpurified TBA in the mobile phases. High background of this ion obscures detection and quantification hydroxybutyric acids. **(C)** Representative extracted ion chromatogram of [103.0401]^-^ in mouse liver metabolite extracts using purified TBA in the mobile phases. Purification removes isobaric inference and enables chromatographic resolution (left to right) of 3-hydroxyisobutyric acid, 3-hydroxybutyric acid, 2-hydroxybutyric acid, and 4-hydroxybutyric acid. Retention time of each isomer was confirmed with analytical standards.

A key strength of TBA ion-pairing chromatography is that it enables retention of highly polar metabolites, including nucleotides, TCA cycle intermediates, and other anionic compounds, on a reversed-phase column. However, many of these analytes are susceptible to adsorption to metal or metal-oxide surfaces in the LC flow path, including stainless steel tubing, frits, column hardware, and other exposed metal components^9–11^. These interactions can impede analyte recovery, broaden peaks, increase carryover, and decrease detected signal. Existing approaches to mitigate these effects include adding medronic acid to the mobile phases and conditioning columns with phosphoric acid. However, medronic acid produces an ion-suppressive background signal during the latter portion of the chromatography, while phosphoric-acid conditioning is labor-intensive and the effect diminishes over time. We therefore sought to optimize metal-mitigation strategies for TBA ion-pairing chromatography.

First, we evaluated whether the concentration of medronic acid could be reduced below the 10 µM concentration previously reported for improving LC–MS analysis of phosphorylated and multicarboxylate metal-sensitive analytes^9^. Lowering medronic acid to 1 µM or 0.1 µM markedly reduced the medronic acid background signal (Fig. 3A). Using ATP as a representative nucleotide, we observed a dose-dependent decrease in ATP peak intensity as medronic acid concentration was reduced (Fig. 3B,C). However, the ATP signal-to-noise ratio was similar between 10 µM and 1 µM medronic acid conditions (Fig. 3D). We therefore proceeded with 1 µM medronic acid. Because the medronic acid signal appeared primarily in the latter portion of the gradient, we reasoned that medronic acid present in mobile phase A may unnecessarily load onto the column during the early aqueous portion of the run and contribute to background when it elutes under higher-organic conditions. Since ATP and other late-eluting metal-sensitive analytes were still exposed to medronic acid as mobile phase B increased, we tested whether medronic acid could be restricted to mobile phase B. This configuration further decreased the medronic acid background signal without reducing ATP intensity (Fig. 3E), indicating that continuous delivery of medronic acid in both mobile phases was not required to preserve nucleotide detection under these conditions.

**Figure 3.**
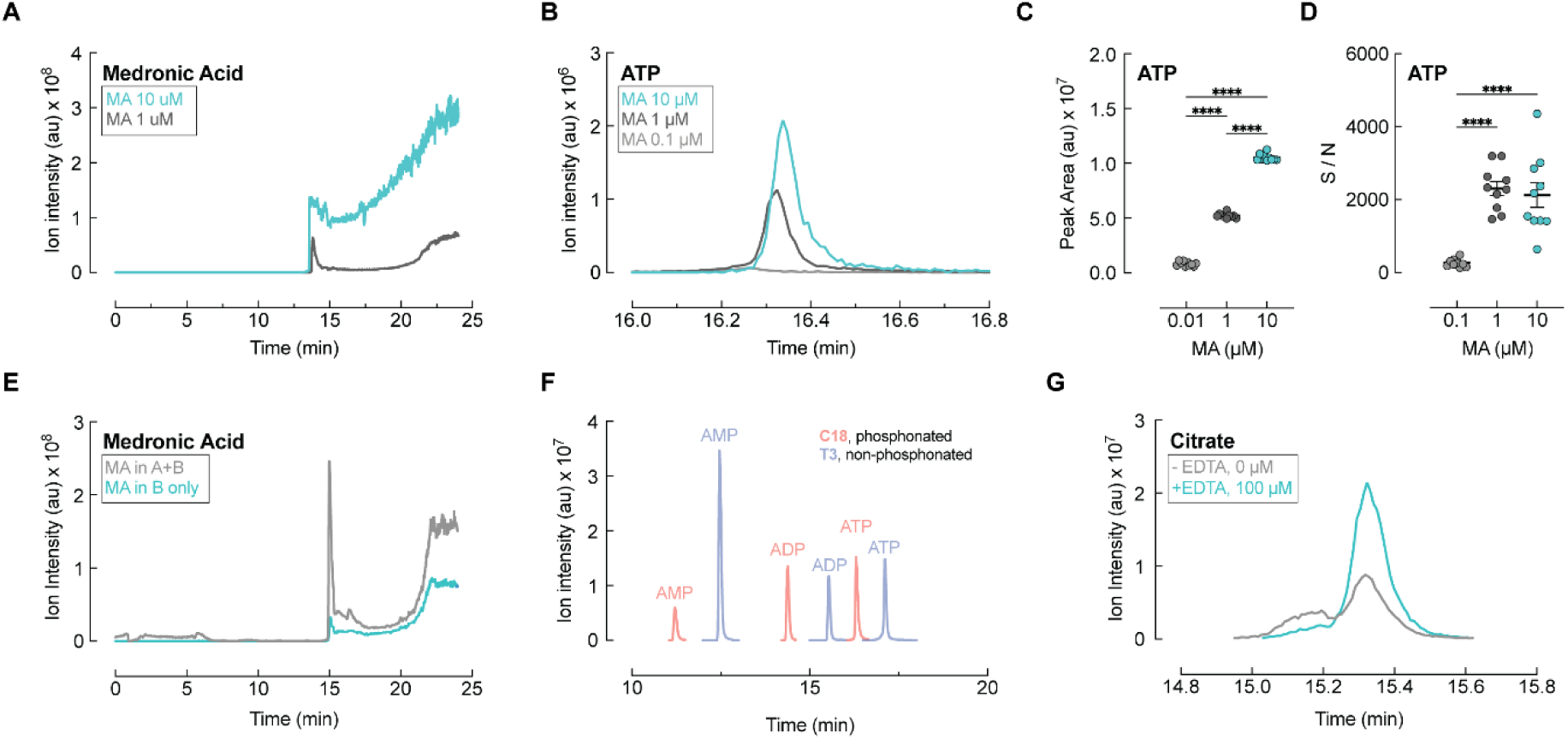
Optimization of cation-interaction mitigation strategies to minimize background and improve ease of use. **(A)** Extracted ion chromatogram of medronic acid [174.9567]^-^ at decreasing from 10 µM to 1 µM and to 0.1 µM. Note that 0.1 µM was not visible on the presented scale and is not included in the plot. Extracted ion chromatogram representative of five independent replicates. **(B)** Effects of mobile phase medronic acid concentration on ATP chromatogram and signal intensity. **(C)** Decreasing mobile phase medronic acid concentration leads to decreased ATP peak area (n=5/group). **(D)** Decreasing medronic acid from 10 to 1 µM preserves the ATP signal to noise (n=5/group). **(E)** Restricting 1 µM medronic acid to mobile phase B only further decreases the background medronic acid signal. Extracted ion chromatogram representative of three independent replicates. **(F)** Both phosphonated C18 and non-phosphonated Acquity Premier T3 columns retain and preserve signal of adenylate nucleotides. Extracted ion chromatogram representative of five independent replicates. **(G)** Addition of 100 µM EDTA to sample resuspension increases citrate intensity and improves peak shape. Extracted ion chromatogram representative of three independent replicates.

In addition to medronic acid, phosphoric-acid conditioning of the LC flow path and column has been used to improve recovery and peak shape for metal-sensitive analytes. In this approach, the column is flushed overnight with 0.5% phosphoric acid in 90:10 acetonitrile:water to condition exposed metal surfaces; prior work showed improved ADP and ATP peak shape, and modestly increased ATP signal, after this treatment ^9^. However, this conditioning step requires extended instrument downtime, careful diversion of phosphate away from the mass spectrometer, and extensive flushing before analysis. To eliminate the need for phosphoric-acid conditioning, we evaluated a metal-passivated bioinert column, the Waters ACQUITY Premier T3, without prior phosphoric-acid treatment. The Premier T3 column performed at least as well as the phosphoric-acid-conditioned C18 column, including for adenylate nucleotides (Fig. 3F), consistent with prior reports that metal-passivated or hybrid-surface LC hardware can reduce adsorption of metal-sensitive acidic analytes^10, 11^.

Finally, citrate remained broad despite reduced medronic acid background and use of metal-passivated column hardware. Because citrate peak shape was not improved by medronic acid, we reasoned that the remaining broadening may reflect solution-phase complexation with residual cations in the injected sample plug rather than adsorption to metal-active surfaces in the LC flow path. To test this, dried extracts were reconstituted in solvent containing 100 µM EDTA. EDTA substantially improved both citrate peak intensity and peak shape (Fig. 3G), consistent with chelation of residual cations before chromatographic separation. Together, these results form a minimized metal- and cation-mitigation strategy for TBA ion-pairing LC–MS by minimizing medronic acid burden, restricting medronic acid to mobile phase B, replacing phosphoric-acid column conditioning with metal-passivated column hardware, and adding EDTA to the sample reconstitution solvent to improve citrate detection.

After establishing conditions that reduce reagent- and additive-derived background while preserving detection of metal-sensitive analytes, we next benchmarked the optimized TBA ion-pairing method against the initial workflow. To focus the comparison on mobile-phase composition, additive burden, and sample-reconstitution changes, both methods were evaluated using the same T3 column (Table 1). Metabolomic analysis of mouse liver extracts showed increased peak area using the optimized method for most compounds annotated on the method (Figure 4A). Bile acids showed one of the most consistent increases across the targeted panel, making them a useful class of metabolites for investigating the basis of improved detection. We therefore focused on a set of late-eluting features at *m/z* 498.2898 that were strongly enhanced optimized conditions.

**Table 1.** Key parameter overview of initial and optimized tributylamine ion pairing chromatography.

| Parameter | Initial method | Optimized method |
| --- | --- | --- |
| TBA source | Unpurified TBA | SPE-purified TBA |
| Column | ACQUITY Premier HSS T3 | ACQUITY Premier HSS T3 |
| Medronic acid | 10 μM, mobile phases A and B | 1 μM, mobile phase B only |
| Sample reconstitution | Water | Water + 100 μM EDTA |
| Gradient | Same gradient | Same gradient |
| MS acquisition | Same acquisition | Same acquisition |

**Figure 4.**
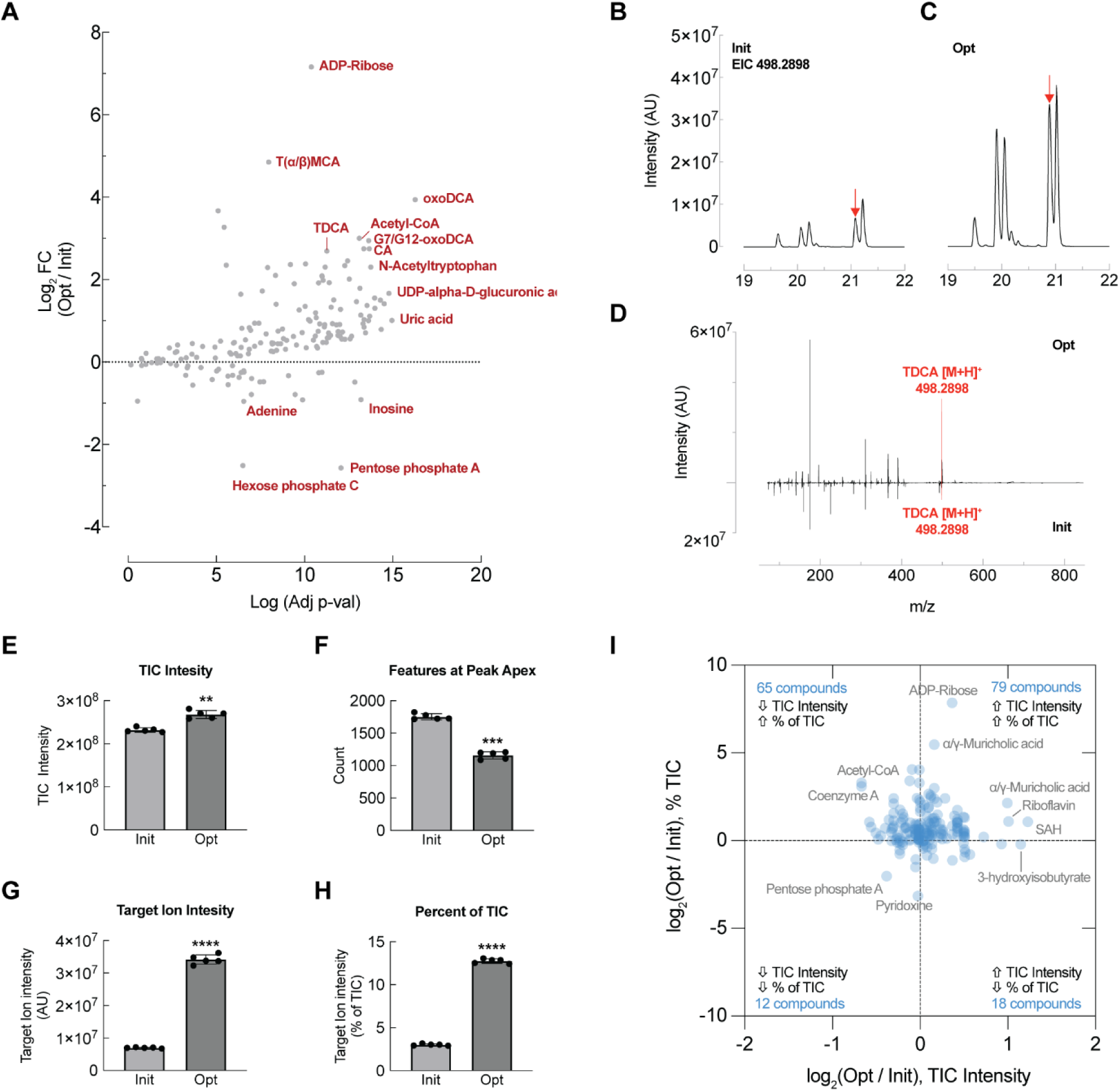
Targeted metabolomics of mouse liver demonstrate broad gain in metabolite intensity in the optimized versus initial TBA IP methods. **(A)** Volcano plot of mouse liver metabolites detected using the optimized (Opt) versus the initial (init) method (n=5 per group). **(B)** Representative extracted ion chromatogram of taurodeoxycholic acid isomers [498.2898]^-^ in the initial method. Representative of n=5 separate injections. **(C)** Representative extracted ion chromatogram of taurodeoxycholic acids [498.2898]- in the optimized method. Ar-rows denote peak apex from which full scan MS1 spectra are presented in panel D. Representative of n=5 separate injections. **(D)** Mirror plot of full scan MS1 spectra of the optimized (top) and initial (bottom) methods taken at the peak apex of the taurodeoxycholic acid peak denoted by the arrows in B and C. Representative of n=5 separate injections. **(E)** The intensity of the TIC at peak apex of the denoted TDCA peak in B-C in the initial and optimized methods (n=5). **(F)** The number of distinct features in the TIC spectra at peak apex of the denoted TDCA peak in B-C in the initial and optimized methods (n=5). **(G)** The target ion intensity (498.2898) at peak apex of the denoted TDCA peak in B-C in the initial and optimized methods (n=5). **(H)** The target ion in-tensity displayed as a percentage of the total ion intensity in the peak apex spectra denoted in B-C in the initial and optimized methods (n=5). **(I)** Scatter plot showing the log2(Opt/Init) fold change of total ion intensity (X) and percent of target ion of spectra (Y) taken at the peak apex spectra of all compounds in A. In panels E-H, statistical significance was determined using paired t-tests and a 5% significance level. **p < 0.001; ***p = 0.0001; ****p < 0.0001.

Five prominent chromatographic peaks were observed at *m/z* 498.2898, corresponding to the deprotonated ion [M−H] of C H NO S with a mass error of +0.67 ppm. This formula is consistent with taurine-conjugated dihydroxy bile acid isomers. Each peak produced MS/MS fragment ions diagnostic of this compound class, including *m/z* 124.0068, corresponding to the taurine anion [C H NO S] ; *m/z* 106.9803, corresponding to taurine anion – NH [C H SO] ; and *m/z* 79.9568, corresponding to the sulfite/sulfur trioxide anion [SO]. Together, the accurate mass, retention behavior, and MS/MS spectra support assignment of these features as isomeric taurine-conjugated bile acids, including taurodeoxycholic acid, taurochenodeoxycholic acid, tauroursodeoxycholic acid, taurohyodeoxycholic acid, and, in mouse liver, tauromurideoxycholic acid. These bile acid signals were markedly higher with the optimized method than with the initial method (Figure 4B–C), an effect that was also evident in full-scan spectra acquired at the chromatographic peak apex (Figure 4D). We therefore used these features to examine whether the improved signal reflected reduced ion suppression, reduced spectral complexity, or a redistribution of the measured ion population toward endogenous metabolites.

Interestingly, the total ion current, or TIC, at the peak apex was higher with the optimized method (Figure 4E), which is not consistent with a simple model in which improved signal results solely from relief of source-level ion suppression by contaminant removal. However, the number of detected features in the peak-apex spectra was substantially lower with the optimized method (Figure 4F). As a result, the bile acid ion of interest had higher absolute intensity (Figure 4G) and represented a larger fraction of the TIC (Figure 4H). Extending this analysis to all compounds annotated by the method, we found that most analytes, 144 of 174, showed an increased fraction of the TIC at peak apex under the optimized conditions.

These data suggest that reducing background ions through TBA purification and lower medronic acid burden decreases spectral complexity and allows a greater fraction of the accumulated ion population to derive from the analyte of interest. This effect may be particularly relevant for ion-trapping Orbitrap instruments, where automatic gain control and maximum injection time determine the ion population accumulated before mass analysis. Conceptually, this resembles the sensitivity gains reported for Orbitrap selected ion monitoring, in which quadrupole filtering increases the fraction of target ions accumulated and improves signal-to-noise for low-intensity metabolites^15^. In the present method, target ions are not preselected; rather, removal of nonbiological background ions appears to increase the useful metabolite fraction of full-scan spectra. TOF instruments do not rely on the same AGC-limited ion-fill process, this global signal-to-noise effect may be less pronounced on TOF platforms, although reduced background should still improve performance by decreasing ion suppression, isobaric interference, spectral complexity, and nonbiological MS/MS acquisition.

We noted an interesting shift in retention time and, in some cases, elution order using the optimized method. For example, late-eluting bile acids eluted approximately 0.1–0.2 min earlier under the optimized conditions (Figure 4B-C). To evaluate this effect more broadly, we analyzed paired liver extracts using both methods and plotted the retention-time change for each annotated analyte ΔRT = RT_optimized_ - RT_initial_ (Figure 5A). Late-eluting compounds, particularly those eluting after approximately 15 min, generally shifted to earlier retention times in the optimized method, whereas earlier-eluting compounds showed more heterogeneous behavior. Notably, some compounds, especially nucleosides and aromatic amino acids, showed increased retention in the optimized method relative to other metabolites that eluted nearby under the initial conditions. We reasoned that these analyte-specific retention-time shifts could alter local co-elution patterns and introduce new ion-suppression relationships, potentially explaining the decreased apparent abundance of some compounds in the optimized method.

**Figure 5.**
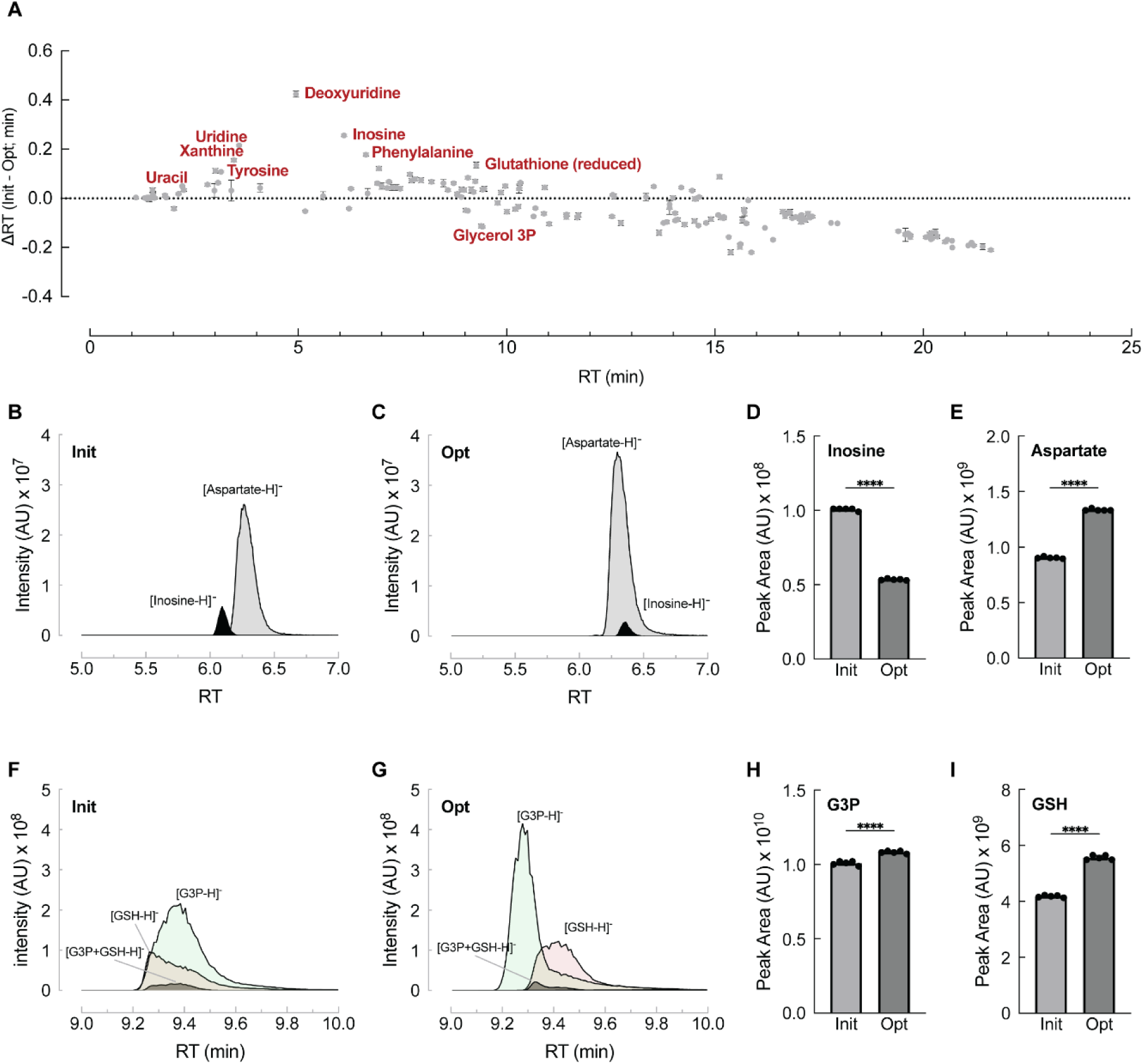
Compound-specific retention-time shifts alter local co-elution and ion suppression. **(A)** Effects of method optimization on compound retention time in mouse liver extracts (n=5). **(B)** Overlaid extracted ion chromatograms of inosine (black) and aspartate (grey) in the initial method. Representative of n=5 replicates**. (C)** Overlaid extracted ion chromatograms of inosine (black) and aspartate (grey) in the optimized method. Representative of n=5 replicates. **(D)** Peak area of inosine in initial and optimized methods (n=5). **(E)** Peak area of aspartate in initial and optimized methods (n=5). **(F)** Overlaid extracted ion chromatograms of G3P (red), GSH (green), and heterodimeric adduct [G3P+GSH-H]- in the initial method. Representative of n=5 replicates. **(G)** Overlaid extracted ion chromatograms of G3P (red), GSH (green), and heterodimeric adduct G3P+GSH-H in the optimized method. Representative of n=5 replicates**. (H)** Peak area of G3P in initial and optimized methods (n=5). **(I)** Peak area of GSH in initial and optimized methods (n=5). In panels D-E and H-I, statistical significance was determined using paired t-tests and a 5% significance level. **p < 0.001; ***p = 0.0001; ****p < 0.0001.

Inosine, a purine nucleoside, was among the compounds with the largest decrease in abundance in the optimized method (Figure 4A). It also eluted approximately 0.25 min later under the optimized conditions (Figure 5A). In contrast, the retention time of the closely eluting compound aspartate was unaffected. Overlaid extracted ion chromatograms showed that inosine and aspartate were chromatographically separated in the initial method (Figure 5B), whereas the specific retention shift of inosine in the optimized method moved it into co-elution with aspartate (Figure 5C). Because aspartate signal is ∼10X that of aspartate, and its signal further increased by ∼ 25% in the optimized method (Figure 5E), this new co-elution likely caused ion suppression of inosine (Figure 5D), reducing its apparent abundance.

Another elution order effect observed was glycerol-3-phosphate (G3P) and reduced glutathione (GSH). These compounds co-eluted under the initial conditions but shifted in opposite directions in the optimized method, with G3P eluting 0.15 min earlier and GSH eluting 0.16 min later (Figure 5A). Extracted ion chromatograms indicated that, in the initial method, co-eluting G3P suppressed the apex of the GSH peak, giving the GSH peak an apparent fronting phenotype Figure 5F-G). Co-elution was also associated with formation of a heterodimeric adduct [G3P+GSH-H]-, further distributing the analyte signal across multiple ion species (Figure 5F). In the optimized method, the differential retention-time shifts led to separation of glycerol-3-phosphate and GSH chromatographically, increasing the observed peak area of both analytes and improving the GSH peak shape (Figure 5G-I). Together, these results show that method optimization can produce analyte-specific changes in retention behavior that reshape local co-elution patterns. In some cases, these shifts improve detection by resolving interfering analytes, while in other cases new co-elution events may decrease apparent abundance through ion suppression.

Finally, we evaluated the performance of the optimized method for untargeted metabolomics across several common sample matrices. Metabolite extracts from pooled mouse liver, GM24385 human B lymphocytes, or NIST SRM 1950 human plasma were analyzed using both the initial and optimized methods. Features were retained for analysis if they were greater than five-fold above the blank and had a peak rating greater than 6 in Compound Discoverer (v3.5). Differential feature abundance analysis showed a broad increase in peak area with the optimized method, with four- to ten-fold more features increasing than decreasing across sample types. In liver extracts, 1,848 features passed filtering, of which 1,385 were significantly increased and 328 were significantly decreased in the optimized method relative to the initial method (Fig. 6A). In B lymphocyte extracts, 1,068 features passed filtering, including 822 increased and 107 decreased features (Fig. 6B). In plasma, 1,188 features passed filtering, of which 1,018 were significantly increased and 103 were significantly decreased (Fig. 6C).

**Figure 6.**
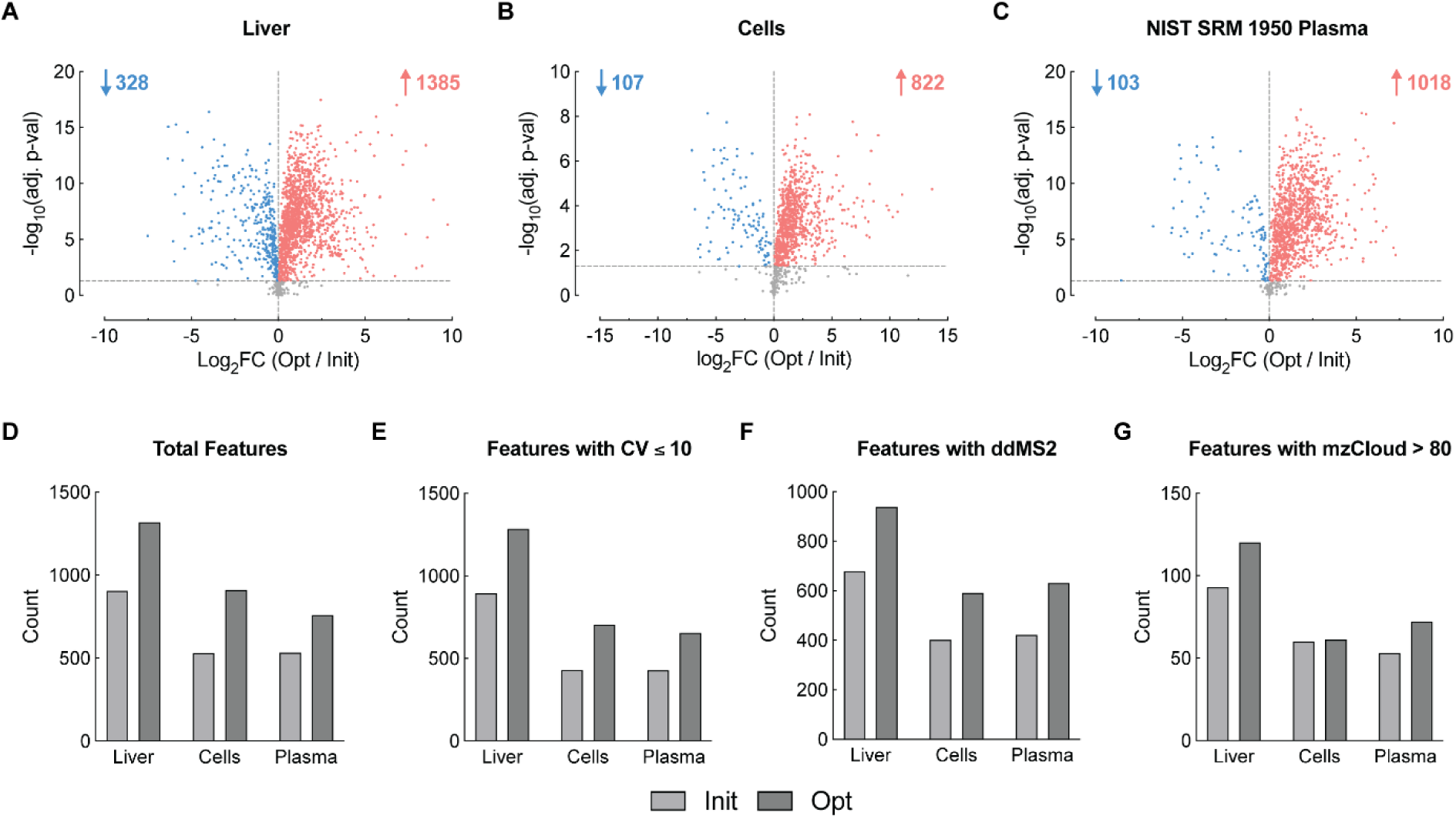
Effects of TBA method optimization on untargeted metabolomics data. **(A)** Untargeted metabolomics of pooled mouse liver tissue analyzed with the initial and optimized methods (n=5). Data were analyzed in Compound Discoverer (v3.5). Presented features were present at >5-fold over blank and had a peak rating of >6 in at least 5 samples. **(B)** Untargeted metabolomics of GM24385 human B lymphocyte analyzed with the initial and optimized methods (n=3). Data were analyzed in Compound Discoverer (v3.5). Presented features were present at >5-fold over blank and had a peak rating of >6 in at least 3 samples. **(C)** Untargeted metabolomics of human SRM1950 plasma analyzed with the initial and optimized methods (n=5). In panels **A-C**, significant features were denoted using an adjusted p-value cutoff of 0.05. Data were analyzed in Compound Discoverer (v3.5). Presented features were present at >5-fold over blank and had a peak rating of >6 in at least 5 samples**. (D)** Total MS1 features with >5-fold over blank peak rating of >6 detected from untargeted metabolomics analysis performed separately on each optimized and initial methods**. (E)** Total MS1 features in (D) with a coefficient of variation (CV) <10% from untargeted metabolomics analysis performed separately on each optimized and initial methods. **(F)** Total MS1 features in (D) with an associated data-dependent MS2 (ddMS2) spectra from untargeted metabolomics analysis performed separately on each optimized and initial methods. **(G)** Total MS1 features in (D) with an mzCloud match score of >80 from untargeted metabolomics analysis performed separately on each optimized and initial methods.

This paired analysis does not fully capture method-specific untargeted performance. When data from the initial and optimized methods are processed together, feature grouping and compound annotation can be driven by signal detected primarily in one method and then propagated across the full dataset. This makes joint processing useful for comparing feature intensity, but less suitable for determining how many features, MS/MS spectra, or annotations each method independently supports.

Separate processing by method showed that the optimized method increased the total number of detected features by 45%, 72%, and 42% in liver, B lymphocytes, and plasma, respectively (Fig. 6D). The optimized method also increased the number of features with variance less than 10% (Fig. 6E), precursor ions with data-dependent MS/MS spectra (ddMS2; Fig. 6F), and MS/MS library matches (Fig. 6G). Collectively, these results indicate that the optimized method improves untargeted metabolomics performance by increasing feature signal intensity and improving acquisition of interpretable MS/MS spectra. This improvement is consistent with reduced background-ion burden, which would otherwise consume data-dependent MS/MS scans on contaminant ions and/or interfere with clean precursor isolation.

Collectively, this work demonstrates that background-ion burden is a major, addressable determinant of TBA ion-pairing LC–MS metabolomics performance. By reducing reagent-derived contaminants, additive background, and metal/cation-associated analyte losses, the optimized workflow improved targeted metabolite signal intensity and increased untargeted feature detection, reproducibility, ddMS2 acquisition, and MS/MS library matching across liver, B lymphocyte, and plasma extracts. These gains are especially important for low-input and single-cell metabolomics, where limited analyte abundance makes nonbiological ion current disproportionately costly. Consistent with this impact, the optimized TBA ion-pairing strategy described here has been adapted to nanoflow LC–MS for low-input polar metabolomics and stable isotope tracing, providing a foundation for single-cell-scale analysis^16, 17^.

More broadly, this study frames background mitigation as a practical design principle for LC–MS method development. The benefits observed here likely reflect multiple mechanisms, including reduced isobaric interference, decreased spectral complexity, improved acquisition of informative MS/MS spectra, and relief of compound-specific electrospray ion suppression. On the Orbitrap platform used here, reduced background may also improve useful ion allocation by increasing the fraction of accumulated ions derived from endogenous metabolites, analogous in concept to the sensitivity gains observed with selected ion monitoring. The magnitude of this ion-budget effect may differ on non-ion-trapping platforms such as TOF instruments, but the broader principle should extend across LC–MS applications: background ions compete with analytes for detection, annotation, and acquisition priority.

In conclusion, improving LC–MS metabolomics is not only a matter of increasing chromatographic retention or instrument sensitivity, but also of increasing the fraction of measured signal that is biologically informative. Background ion burden is itself an optimizable determinant of method performance. Systematic identification and mitigation of reagent-, additive-, hardware-, and sample-derived background provides a generalizable route to cleaner spectra, higher useful signal, improved MS/MS acquisition, and greater annotation confidence. For TBA ion-pairing metabolomics, this approach preserves the broad coverage and retention stability of reversed-phase ion-pairing while making the method more sensitive, interpretable, and adaptable to sample-limited applications.

## Acknowledgments.

LC-MS analyses were conducted in the Van Andel Institute Mass Spectrometry Core (RRID:SCR_024903). This work was financially supported by the Van Andel Institute MeNu (<u>Me</u>tabolism and <u>Nu</u>trition) program (RRID: SCR_027494). A.A.C. is the owner of Caudy Bioanalytical Advisory. The author received no payment for this work and declares no specific financial competing interest related to the manuscript. The other authors have no conflicts of interest. The authors thank Dr. Adam Rosebrock for many helpful and insightful discussions on these topics.

## Materials and methods

### Biological samples and experimental design

Mouse liver, GM24385 human B lymphocytes, and NIST SRM 1950 human plasma were used to evaluate method performance across biological matrices. Pooled mouse liver extracts were used for method-optimization experiments and for comparison of initial and optimized LC–MS conditions. GM24385 B lymphocytes and NIST SRM 1950 plasma were used to evaluate untargeted metabolomics performance across sample types. Five technical replicates of mouse liver and plasma were analyzed for matrix comparisons. For B lymphocyte analyses, three technical replicates from each of four drug-treated groups were analyzed, and sample injection order was randomized. Samples were stored at −80 °C before analysis. Water blanks and process blanks were included in each analysis.

### Metabolite extraction from liver tissue and suspension cells

Polar metabolites from liver tissue and suspension cells were extracted using a modified Bligh-Dyer extraction^18, 19^ with a final chloroform:methanol:water ratio of 2:2:1.8 v/v/v. Liver tissue was homogenized in ice-cold 1:1 chloroform:methanol using chloroform (Honeywell, 650471-1L) and LC–MS-grade methanol (Fisher Scientific, A456-4) at a ratio of 40 mg tissue per 1 mL extraction solvent. Suspension cells were extracted at a ratio of 2 × 10 cells per 1 mL extraction solvent.

After solvent addition, samples were vortexed for 10 s, homogenized using a liquid nitrogen-chilled bead mill for 30 s at 6 m/s, sonicated in a water bath for 5 min, and incubated on wet ice for 30 min. LC–MS-grade water (Fisher Scientific, W6-4) was then added to achieve the final 2:2:1.8 chloroform:methanol:water ratio. For liver tissue, sample water content was assumed to be 70% and was accounted for when calculating the water addition. Samples were vortexed for 10 s, incubated on wet ice for 10 min, and centrifuged for 10 min at 17,000 × g and 4 °C to separate the lower organic phase, upper polar phase, and interphase protein pellet.

For liver samples, an aliquot corresponding to 34 mg tissue equivalents was collected from the upper polar phase and centrifuged a second time to remove residual particulates. A clarified aliquot corresponding to 32 mg tissue equivalents was dried under vacuum and reconstituted in 200 µL LC–MS-grade water. In some experiments as indicated in text, samples were reconstituted in 100 µM EDTA (Thermo Fisher Scientific, AM9260G). For suspension cell samples, an aliquot corresponding to 1.6 × 10 cell equivalents was collected from the upper polar phase, dried under vacuum, and reconstituted in 50 µL LC–MS-grade water containing 100 µM EDTA. For initial method comparisons, EDTA was omitted from the reconstitution solvent. L-glutamic acid-2,3,3,4,4-d (Cambridge Isotope Laboratories, DLM-556-PK) was added to all samples as an internal standard at a final concentration of 1 µg/mL. Reconstituted samples were transferred to autosampler vials with inserts (Sigma-Aldrich, 29391-U) before LC–MS analysis.

### Metabolite extraction from plasma

Polar metabolites from plasma were extracted using ice-cold acetonitrile:methanol:water, 4:4:2 v/v/v, prepared with LC–MS-grade acetonitrile (Fisher Scientific, A955-4), methanol (Fisher Scientific, A456-4), and water (Fisher Scientific, W6-4)^19^. Plasma was extracted at a ratio of 40 µL plasma per 1 mL extraction solvent. Samples were vortexed for 10 s, sonicated in a water bath for 5 min, incubated on wet ice for 1 h, and centrifuged for 10 min at 17,000 × g and 4 °C. An aliquot of supernatant corresponding to 32 µL plasma equivalents was dried under vacuum and reconstituted in 100 µL LC–MS-grade water containing 100 µM EDTA. For initial method comparisons, EDTA was omitted from the reconstitution solvent. L-glutamic acid-2,3,3,4,4-d was added as an internal standard at a final concentration of 1 µg/mL.

### Solid-phase purification of tributylamine

Tributylamine (TBA; Sigma-Aldrich, 90780-100ML) was purified by serial solid-phase extraction using polymeric reversed-phase, strong anion-exchange, and strong cation-exchange sorbents. Strata-X polymeric reversed-phase cartridges (Phenomenex, 8B-S100-JEG), Strata-X-A polymeric strong anion-exchange cartridges (Phenomenex, 8B-S123-JEG), and Strata-X-C polymeric strong cation-exchange cartridges (Phenomenex, 8B-S029-JEG) were mounted on a 12-port vacuum SPE manifold (Phenomenex, VM12). Each cartridge contained a 1 g sorbent bed in a 20 mL cartridge format.

Cartridges were washed with 5 mL LC–MS-grade methanol followed by two washes with 5 mL LC–MS-grade water and were allowed to drain completely between washes. Each cartridge was then conditioned with 2 mL TBA. Approximately 80 mL TBA was passed sequentially through Strata-X, Strata-X-A, and Strata-X-C cartridges at approximately −10 inHg and ∼5 mL/min. The flow-through was collected in ∼10 mL aliquots in glass conical tubes (PYREX; Sigma-Aldrich, CLS995025-125EA), pooled, and used to prepare mobile phases for the optimized LC–MS method. Glass serological pipettes (VWR, 93000-698) were used for TBA handling.

### Elution of TBA-derived contaminants from SPE cartridges

To profile contaminants captured during TBA purification, Strata-X, Strata-X-A, and Strata-X-C cartridges were prepared as described above. A 10 mL aliquot of TBA was passed through individual cartridges independently. Matched process blanks were prepared using LC–MS-grade water in place of TBA. After loading, retained compounds were eluted from each cartridge using two 5 mL aliquots of elution solvent. Strata-X cartridges were eluted with 2% formic acid (Fisher Scientific, A117-50) in 1:1 methanol:acetonitrile. Strata-X-A cartridges were eluted with 5% formic acid in methanol. Strata-X-C cartridges were eluted with 5% ammonium hydroxide (Fisher Scientific, A470-250) in methanol. A 1 mL aliquot of each eluate was dried under vacuum and reconstituted in 100 µL LC–MS-grade water before LC–MS analysis. Three technical replicates of each TBA eluate and two technical replicates of each process blank were analyzed.

### Mobile phase preparation

A TBA buffer additive was prepared by combining 900 mL LC–MS-grade methanol, 71.5 mL TBA, and 25.75 mL LC–MS-grade acetic acid (Fisher Scientific, A113-50). The additive was mixed thoroughly before use. Unless otherwise indicated, mobile phase A consisted of LC–MS-grade water containing 3% LC–MS-grade methanol, 10 mM TBA, and 15 mM acetic acid. Mobile phase B consisted of LC–MS-grade methanol containing 10 mM TBA, 15 mM acetic acid, and medronic acid (Agilent Technologies, 5191-4506). In the optimized method, SPE-purified TBA was used for both mobile phases and 1 µM medronic acid was included only in mobile phase B. Mobile phases were mixed thoroughly and sonicated before use. The column-wash mobile phase consisted of 99:1 acetonitrile:water.

### Initial and optimized LC–MS method conditions

Method performance was compared between an initial TBA ion-pairing workflow and the optimized workflow. The optimized method incorporated SPE-purified TBA, 1 µM medronic acid restricted to mobile phase B, EDTA in the sample reconstitution solvent, and a metal-passivated column. The initial workflow used unpurified TBA, medronic acid in both mobile phases, and no EDTA in the reconstitution solvent. Although the original initial method used a phosphoric-acid-conditioned ZORBAX RRHD Extend-C18 column, 2.1 × 150 mm, 1.8 µm (Agilent, 759700-902), with a ZORBAX Extend-C18 UHPLC guard cartridge (Agilent, 821725-907), direct comparisons of initial and optimized mobile-phase conditions were performed using the same ACQUITY Premier HSS T3 column configuration to isolate the effects of reagent and mobile-phase optimization.

### Phosphoric-acid conditioning of C18 columns

Where indicated, C18 columns were conditioned with phosphoric acid before use. A conditioning solution was prepared by combining 540 mL LC–MS-grade acetonitrile, 60 mL LC–MS-grade water, and 3 mL of 85% phosphoric acid (Sigma-Aldrich, 49685-100ML), corresponding to 0.5% phosphoric acid in 90:10 acetonitrile:water. LC lines were first purged with LC–MS-grade water at 5 mL/min for 5 min, and the flow path and column were flushed with LC–MS-grade water at 0.25 mL/min for 30 min. LC lines were then transferred to the phosphoric-acid conditioning solution and purged at 5 mL/min for 5 min. The column was conditioned overnight at 0.1 mL/min for at least 12 h.

After conditioning, LC lines were returned to LC–MS-grade water and purged at 5 mL/min for 15 min. The column was disconnected, and eluent pH at the column inlet was measured at 0.1 mL/min. If the pH was below 4.5, the flow path was washed with water until the pH exceeded 4.5. The column was then reconnected and flushed with LC–MS-grade water at 0.25 mL/min for 1 h. During the entire process, LC lines were fully disconnected from the mass spectrometer.

### LC–MS analysis

Samples were analyzed using a Vanquish binary pump chromatography system coupled to an Orbitrap Exploris 240 mass spectrometer equipped with a heated electrospray ionization source (Thermo Fisher Scientific). Data were acquired in negative-ion mode. Unless otherwise indicated, 2 µL of each sample, blank, or standard was injected. Chromatographic separation was performed using an ACQUITY Premier HSS T3 column, 2.1 × 150 mm, 1.8 µm, with VanGuard FIT hardware (Waters, 186009472). The column temperature was maintained at 35 °C, and the flow rate was 0.25 mL/min. The analytical gradient was 24 min: 0–2.5 min, 0% B; 2.5–7.5 min, 0–20% B; 7.5–13 min, 20–45% B; 13–20 min, 45–99% B; and 20–24 min, 99% B. A 16 min reverse-flow wash and re-equilibration method was run between injections. During this method, the wash mobile phase B was 99:1 acetonitrile:water. The wash gradient was: 0–3 min, 100% B at 0.25 mL/min; 3–3.5 min, 100% B with flow ramped to 0.8 mL/min; 3.5–7.35 min, 100% B at 0.8 mL/min; 7.35–7.5 min, 100% B with flow ramped to 0.6 mL/min; 7.5–8.25 min, 100–0% B with flow ramped to 0.4 mL/min; 8.25–15.5 min, 0% B with flow ramped to 0.25 mL/min; and 15.5–16 min, 0% B at 0.25 mL/min. Electrospray source parameters were: spray voltage, −2500 V; sheath gas, 60 arbitrary units; auxiliary gas, 19 arbitrary units; sweep gas, 1 arbitrary unit; ion transfer tube temperature, 320 °C; and vaporizer temperature, 250 °C. Full-scan MS1 data were acquired over *m/z* 70–850 at 240,000 resolution, with the RF lens set to 35%. Data-dependent MS/MS spectra were acquired using higher-energy collisional dissociation with stepped collision energies of 15, 30, and 45. MS/MS spectra were acquired at 15,000 resolution with a 2 *m/z* isolation window and a maximum of five data-dependent scans per duty cycle. Targeted MS/MS inclusion triggers were used where indicated.

System suitability was assessed by injecting a standard mixture before and after each analytical batch. The mixture contained L-glutamic acid, L-histidine hydrochloride, orotic acid, ATP, and malate at 1 µg/mL each and was stored at −80 °C. Standards were prepared from L-glutamic acid and L-histidine hydrochloride from the Supelco amino acid standard set (Supelco, LAA21-1KT), orotic acid (Sigma-Aldrich, O2750-10G), adenosine 5′-triphosphate disodium salt hydrate (Sig-ma-Aldrich, A2383-1G), and L-(−)-malic acid (Sigma-Aldrich, M7397-25G).

### Medronic acid and EDTA optimization

To determine whether medronic acid concentration could be reduced while preserving nucleotide performance, mobile phases were prepared with medronic acid at 10 µM, 1 µM, or 0.1 µM. ATP was used as a representative nucleotide to evaluate peak intensity, peak shape, and signal-to-noise ratio. Additional experiments tested whether 1 µM medronic acid could be restricted to mobile phase B rather than included in both mobile phases. Medronic acid background signal and ATP performance were compared across conditions.

To evaluate whether citrate peak shape was affected by residual cations in the injected sample, dried metabolite extracts were reconstituted with or without 100 µM EDTA in LC–MS-grade water. Citrate peak intensity and peak shape were compared between conditions. The optimized method used 100 µM EDTA in the sample reconstitution solvent.

### Data processing, visualization, and statistical analysis

Untargeted data analysis was performed in Compound Discoverer v3.5 (Thermo Fisher Scientific). Raw LC–MS files from TBA SPE eluates, matched process blanks, and biological matrix extracts were processed for chromatographic alignment, feature detection, compound grouping, blank filtering, and putative annotation. For TBA-contaminant eluate analysis, features were retained if they had a peak rating >7, intensity >1 × 10, and were enriched at least 10-fold relative to matched water-process blanks. For untargeted analysis of mouse liver, human B lymphocyte, and human plasma extracts, features were retained if they were >5-fold above blank and had a peak rating >6. Differential feature-intensity analysis was performed between the initial and optimized methods, and features with adjusted *p* < 0.05 were considered significant.

For method-specific feature-count comparisons, initial and optimized datasets were processed separately in Compound Discoverer to determine the number of detected MS1 features, features with coefficient of variation <10%, features with associated data-dependent MS/MS spectra, and features with mzCloud library match scores >80. Extracted ion chromatograms and representative full-scan or MS/MS spectra were exported from Thermo Freestyle and plotted in GraphPad Prism. Statistical analyses and figure generation were performed in GraphPad Prism. Sample sizes are indicated in the corresponding figure legends.

